# Expanding the options for therapeutic exon skipping as a future treatment for *USH2A*-associated disease by 3D structural modeling of newly formed hybrid domains

**DOI:** 10.64898/2026.04.24.720583

**Authors:** Lucija Malinar, Sanne Broekman, Daniel T. Rademaker, Anh Q. Le, Theo Peters, Erik de Vrieze, Peter A.C. ‘t Hoen, Erwin van Wijk, Hanka Venselaar

**Affiliations:** Department of Medical BioSciences, Radboud University Medical Center, Geert Grooteplein Zuid 26, 6525 GA Nijmegen, Netherlands; Eye Hospital, University Medical Centre Ljubljana, 1000 Ljubljana, Slovenia; Amsterdam Machine Learning Lab, University of Amsterdam, Science Park 900, 1098 XH Amsterdam, The Netherlands; HIMS-Biocat, University of Amsterdam, Science Park 904, Amsterdam 1098 XH, The Netherlands; Biosystems Data Analysis, University of Amsterdam, 1090 GE Amsterdam, the Netherlands; Biomedical Sciences, Leiden University, Hippocratespad 21, 2333 ZD Leiden, the Netherlands; Department of Otorhinolaryngology, Hearing and Genes, Radboud University Medical Center, Geert Grooteplein 10, 6525 GA Nijmegen, the Netherlands

**Keywords:** Exon skipping, 3d modelling, bioinformatics, protein struture, USH2a

## Abstract

Usher syndrome, the leading cause of hereditary deaf-blindness affecting approximately 1 in 15,000 individuals worldwide, is currently still untreatable. Antisense oligonucleotide-based exon skipping has shown significant therapeutic promise for *USH2A*-associated retinal dysfunction. Selection of (combinations of) exons suitable for therapeutic exon skipping within the fibronectin type 3 (FN3) domain-encoding region of *USH2A* currently requires that skipped exons exactly align with complete protein domains. However, only few exon combinations meet this criterion, which significantly restricts the therapeutic potential of this strategy. Our study addresses this limitation by incorporating AlphaFold2 structural modelling into the exon skipping target selection pipeline. Following this adjusted framework, we can predict exon skipping combinations that allow remaining domain fragments to form structurally viable hybrid domains. As a proof-of-concept, we examined and confirmed the functionality of usherinΔexon54-58 that contains a hybrid FN3 domain, using zebrafish as a model. This highligts the potential of the newly developed paradigm for identifying exon skipping targets with potential therapeutic relevance. Our results emphasize the value of structural modeling in identifying new therapeutic exon skipping targets, aiming to improve precision, efficiency, applicability, and cost-effectiveness in the development of genetic therapies for hereditary diseases such as Usher syndrome.

## INTRODUCTION

Usher syndrome, affecting an estimated 1 in 15,000 individuals worldwide, is the primary cause of combined hereditary sensorineural hearing and vision loss in man (1). It is genetically heterogeneous, with pathogenic variants in 10 different genes implicated in its manifestation. Among these, the *USH2A* gene is the most frequently mutated gene, explaining ∼50% of all Usher syndrome cases and up to 80% of individuals diagnosed with Usher syndrome type 2 (USH2) (2). Besides that, pathogenic variants in *USH2A* are also the most frequent cause of autosomal recessively inherited non-syndromic retinitis pigmentosa (arRP), explaining up to 12-25% of cases (3). To date, 2764 unique variants of *USH2A* gene have been listed in the LOVD database (visited on the 12th of February 2026) (4), with 1760 classified as being (likely) pathogenic. While currently no treatment options exist for the Usher syndrome-associated vision loss (5), the experienced hearing loss can be alleviated through the use of hearing aids and cochlear implants. The severe and debilitating aspects of Usher syndrome impose a significant burden on affected individuals and society as a whole, emphasizing the urgency for developing effective treatments.

Exon skipping using antisense oligonucleotides (ASOs) has emerged as a promising therapeutic approach, offering an alternative for gene augmentation to tackle diseases caused by pathogenic variants in genes that encode large structural proteins with a repetitive domain architecture, such as Duchenne Muscular Dystrophy (dystrophin), *USH2A*-associated retinal disease (usherin), and CADASIL (NOTCH3) (6–8). This method is particularly useful in conditions where gene augmentation strategies face challenges related to the limited packaging capacity of the currently used viral vehicles for gene delivery and the presence of multiple protein isoforms. ASOs are designed (9–11) to induce the targeted in-frame skipping of (frequently) mutated endogenous exons during the process of pre-mRNA splicing. In many cases, it allows for the production of a modified, yet functional protein, circumventing the detrimental effects of pathogenic genetic variants. As the technique can be customized to suit the specific genetic profiles of individual patients, it offers both the potential to make significant advancements in the development of personalized treatments and the broader effort to overcome genetic disorders. However, not all genes are equally amenable to exon skipping-based approaches. An example of this contrast lies in the structural and functional differences between the proteins involved in Duchenne Muscular Dystrophy (DMD) and Usher syndrome type 2A (USH2A).

In DMD, most cases are caused by genomic rearrangements in the dystrophin gene (*DMD*) that disrupt the open reading frame. Targeted exon skipping can restore the reading frame, enabling the production of a shorter, yet partially functional dystrophin protein. The success of this approach is largely attributed to the modular and flexible architecture of dystrophin’s central rod domain, which is composed of spectrin-like (also called pectin) repeats. This gene tolerates internal in-frame deletions remarkably well. Up to 36 exons, that collectively encode the major part of the central rod domain and roughly 46% of the total protein, can be missed without completely abolishing dystrophin’s function (12). This structural flexibility allows many exon combinations in the middle region of the gene to be skipped, while still preserving essential functions at the N- and C-terminal region of dystrophin. As a result, the severe DMD phenotype can be shifted to the milder Becker muscular dystrophy.

In contrast, exon skipping strategies for *USH2A*, which encodes usherin, face significantly more constraints due to the rigid and highly structured domain architecture of the protein. Usherin contains multiple fibronectin type III (FN3) domains, which rely on tightly conserved 3D folding for proper function. Unlike the flexible spectrin repeats in dystrophin, FN3 domains are not modular in the same way: their function depends on the precise positioning of beta-strands and conserved core residues. Any truncation or deletion within an FN3 domain comes at the risk of disrupting its folding, leading to destabilization, and ultimately loss of protein function. As a result, usherin is much less tolerant to random internal deletions, and indiscriminate skipping of in-frame exons will therefore likely not result in the production of shortened usherin protein with sufficient residual function.

The current *USH2A* exon skipping strategies targeting the FN3 domain-encoding region aim to remove entire domains in which pathogenic variants have been identified. This approach requires careful selection of exon combinations that encode complete domains to avoid disrupting (neighboring) domains, adding complexity and limitations to therapeutic exon skipping designs. For example, Schellens *et al*. (13) successfully excised *ush2a* exons 30–31 and 39–40 from the zebrafish genome using CRISPR-Cas9, each of which resulted in the deletion of a single FN3 domain, without compromising usherin expression and subcellular localization. Alternatively, Dulla *et al*. (14) induced the skipping of *USH2A* exon 13, resulting in the removal of three complete and two partial EGF-like domains, which led to the formation of a hybrid EGF-like domain that retained functional properties, both in zebrafish models and in a first-in-man study, despite its structural modificatons.

In the current study, two hypotheses are proposed to advance exon-skipping strategies for regions encoding usherin FN3 domains. First, we propose to generate functional hybrid FN3 domains through selective exon skipping as previously demonstrated for EGF-like domains. Second, we propose to use 3D protein modeling to predict and assess the structural consequences of specific exon-skipping combinations. As a case study, we examined the excision of *USH2A* exons 54–58, which results in a predicted hybrid fibronectin type III (FN3) domain. Based on our *in silico* models in both human and zebrafish, we demonstrated *in vivo* that skipping of exons 54-58 results in a shortened and functional usherinΔexon54–58 protein in which key structural elements are preserved, supporting the predictive value of structure-guided exon skipping design.

## RESULTS

### 3D structural modeling outperforms sequence-based 2D modeling in defining domain boundaries

Determining protein domain boundaries is typically done using resources such as UniProt and SMART (16). These tools enable the mapping of protein domains based on amino acid sequences (2D modeling) to their corresponding exons, providing key insights into the protein’s domain architecture. However, due to the lower conservation of inter-domain regions compared to the domains themselves, these tools often fail to identify the precise structural boundaries of the domains. This uncertainty is directly reflected in the inconsistencies observed between UniProt and SMART predictions in the total number of and identified domain boundaries for FN3 domains in the usherin protein (**Figure 1**). SMART predicts 32 FN3 domains with an average length of 83 amino acids, while UniProt predicts 34 FN3 domains that consist of 98 amino acids on average, resulting in an average 15.3% difference in length. Additionally, UniProt fails to predict a domain in the region spanning amino acids 3,209 to 3,403 (**Figure 1A**), while SMART (**Figure 1B**) predicts an additional FN3 domain spanning amino acid 3,110 to 3,485 (376 amino acids), which by far exceeds the average length of the other FN3 domains. AlphaFold2 3D structure predictions, on the other hand, provide a clear visualization of the domain boundaries, as shown in **Figure 1C**. AlphaFold2 predicts a total of 34 FN3 domains and an additional large, previously unknown ‘flower-like’-protrusion ranging from amino acids 3,209 to 3,403 (‘flower-like’ domain, **Figure 1C**). This demonstrates that combining sequence-based tools like UniProt and SMART with structure-based prediction tools like AlphaFold2 allows for a more comprehensive understanding of the overall 3D structure of a protein and domain boundaries, which is particularly important in the context of developing exon skipping-based therapeutic interventions.

**Figure 1.**
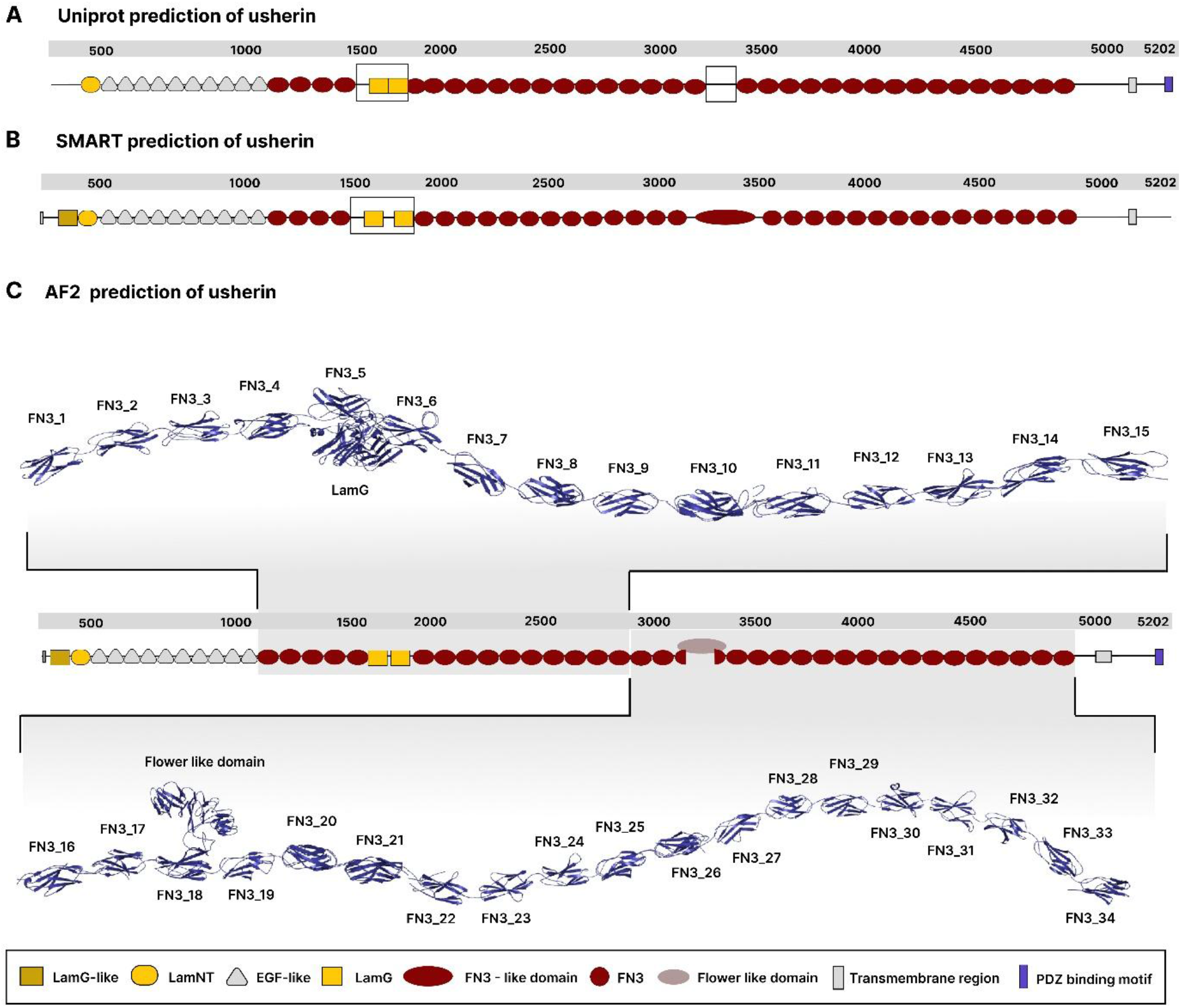
Schematic representation of human usherin based on different prediction algorithms. **A)** 2D representation of usherin based on UniProt prediction data, highlighting consistent domain boundaries. Regions that show gaps are annotated with black boxes. **B)** 2D representation of usherin according to SMART predictions with black boxes indicating potential domain boundaries disputes. **C)** Representation of the most likely usherin protein structural model based on 3D structural models predicted by AlphaFold2.

### FN3 consensus model

FN3 domains require a tightly conserved 3D folding for function, with essential beta-strands and conserved core residues that maintain the domain’s structural integrity. Our consensus model of the FN3 domains in usherin (**Figure 2**) highlights both the conserved and variable regions across these domains. While the core beta-strands (β1-β8) maintain a relatively stable conformation with low flexibility, the loop regions connecting these strands show notable variation. Some loops are longer and differently oriented in space, which can lead to apparent structural differences between FN3 domains. To quantify these spatial variations, a TM-align matrix was used to calculate the TM-score for each FN3 domain (**Supplementary Figure 1**). The TM-score is a normalized measure of structural similarity between two protein folds, ranging from 0 to 1, where values above ∼0.5 typically indicate the same fold and values below ∼0.3 suggest unrelated structures. In our dataset, a pairwise FN3 domain comparison resulted in an average TM-score of 0.81 (ranging from 0.53 to 0.97), indicating that these FN3 domains, with the exception of FN18 and FN19, share a high degree of structural similarity. By placing more weight on the conserved β-strands and less on the flexible loops, TM-score captures the inherent loop flexibility while preserving the structural consistency of the core β-strand framework.

**Figure 2.**
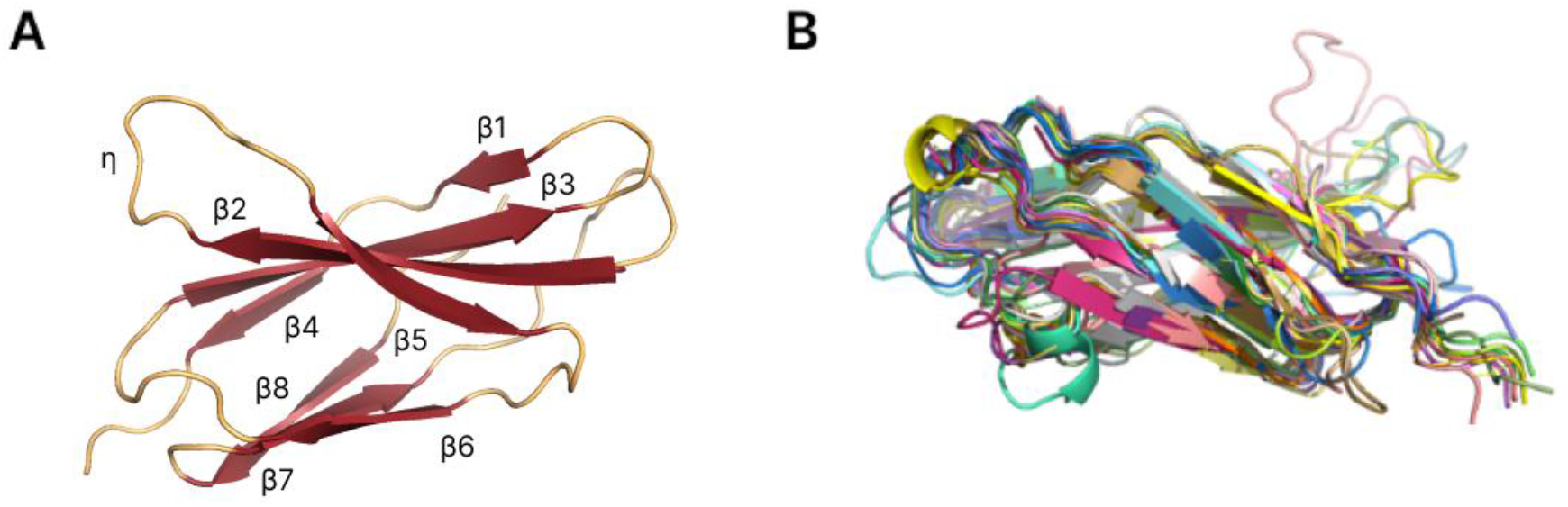
Structural alignment and variability of usherin FN3 domains. **A)** A 3D representation of a single FN3 domain within usherin, highlighting the conserved β-strand core (β1–β8) and a single helix (η). The β-strands are labeled and depicted in yellow, with loops and connecting regions in red, illustrating the stable core structure and flexible loop regions. **B)** An overlay of multiple FN3 domains from usherin, each shown in a different color to emphasize structural similarities and variations. The alignment highlights the conserved folding of the β-strands while revealing significant variability in the loop regions. This structural flexibility suggests potential functional adaptations while maintaining the stability of the core FN3 fold.

### Expanding the therapeutic exon skipping options for *USH2A* using the hybrid domain approach: a case study of skipping exons 54-58

Traditional exon skipping strategies rely on *a priori* selection of in frame exons, that harbor (multiple) recurrent pathogenic variants identified in patients, and that do not interfere with the overall protein domain architecture upon skipping, based on sequence-based prediction tools such as SMART and UniProt. Building on previous research that identified specific in frame exons and associated pathogenic variants using the LOVD database (4), our investigation aimed to explore new therapeutic targets by leveraging the formation of hybrid domains. Initial success was achieved by skipping single, complete FN3 domains in usherin (13), demonstrating the feasibility of exon skipping for therapeutic purposes, while maintaining functional protein structure. Taking this approach further, we investigated the potential of skipping multiple exons to address regions with clusters of pathogenic variants. Following this approach we identified exons 54-58, that collectively encode multiple FN3 domains and in which 188 different (likely) pathogenic variants have been identified. *USH2A* exons 54-58 encode two-thirds of FN3_19, complete FN3_20 and FN3_21, and one-third of FN3_22. *In silico* modeling of the effect of skipping human *USH2A* exons 54-58 using AlphaFold2 (**Figure 3B**) confirmed the 2D schematic representation (**Figure 3A**) indicating that skipping of *USH2A* exons 54-58 results in the formation of a hybrid FN3_19-22 domain. Hybrid domains were first evaluated using the Root Mean Square Deviation (RMSD), with values below 2 Å indicating high structural similarity, 2–3 Å suggesting moderate deviations, and values above 3 Å implying potential misfolding and functional loss. Comparative structural analysis revealed that the newly formed hybrid FN3_19-22 domain closely resembles both the parental FN3_19 (RMSD 0.942) and FN3_22 (RMSD 1.67) domains (**Figure 3C, D**). The RMSD of hybrid and FN3_22 obtains a higher score than that of hybrid and FN3_19, which can be expained by the fact that the *strand*-*β* and *loop*-*α* from the hybrid are not perfectly aligned with same region in the FN3_22. To put it in broader perspective, FN3_19-22 hybrid was structurally aligned to all FN3 domains, and perfectly fitted into the consensus FN3 model (**Figure 3E**). In zebrafish, skipping of *ush2a* exons 54-58 is predicted to result in similar structural outcomes. The newly formed hybrid FN3_19-22 domain in zebrafish usherin resembles the parental FN3 domains 19 (RMSD 2.323 Å) and 22 (RMSD 0.423 Å), with FN3_19 giving the higher score due to longer beta strand *γ* in the hybrid (**Figure 3F, G**). We also calculated the TM-align scores for the newly formed human hybrid compared to the parent domains FN3_19 and FN3_22 as 0.43 and 0.69 respectively. This latter score seems promising since it indicates a strong structure similarity with FN22. Addtionally, the comparison of the human hybrid domain with all other human FN3 domains resulted in scores ranging between 0.58 and 0.86 (excluding domain FN18 which contains the ‘flower-like’ protrusion). (**Supplementary Figure 1**) This strengthens our conclusion that the 3D-structure of hybrid FN19-22 shows sufficient resemblence to the standard FN3 domains.

**Figure 3.**
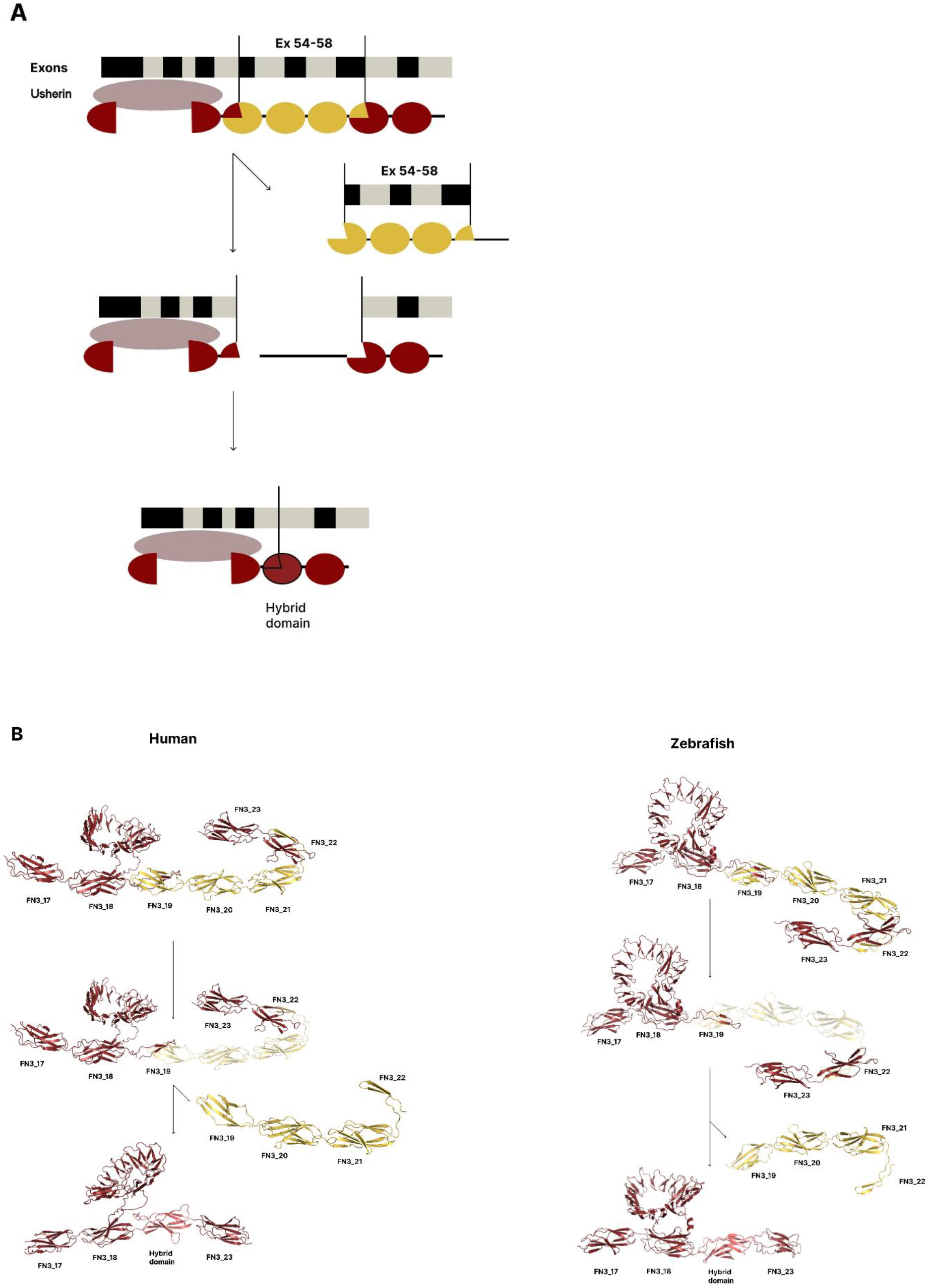
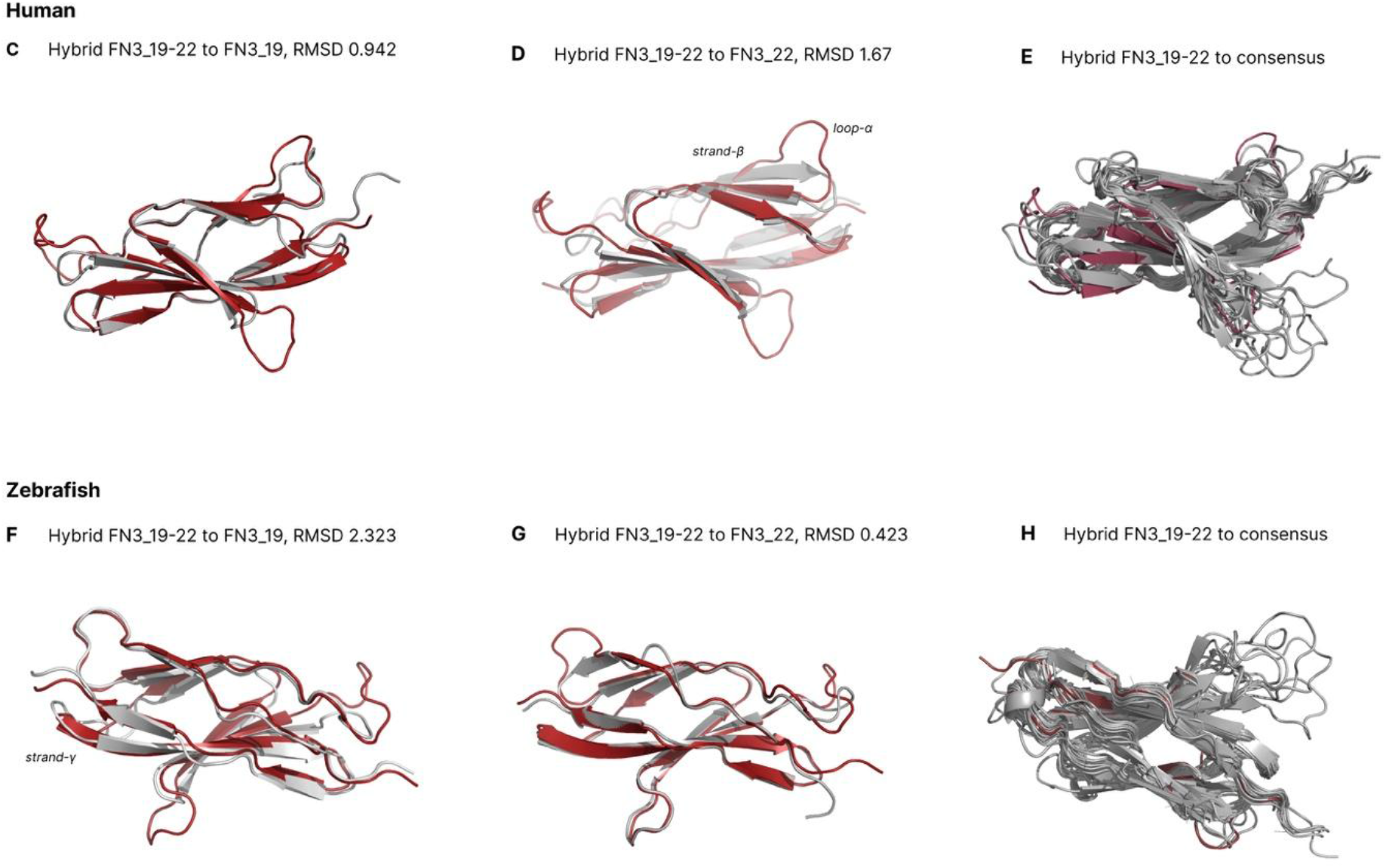
2D Schematic representation and 3D AlphaFold2 models of exon 54-58 skipping in human and zebrafish usherin: **A)** 2D schematic representation of the effects of *USH2A* exon 54-58 skipping at the usherin protein level. The diagrams illustrate the structural outcomes of exon skipping on the FN3 domains 17-23. The segments to be skipped are highlighted in yellow, encompassing two-thirds of FN3_19, complete FN3_20 and FN3_21, and one-third of FN3_22. Skipping of exon 54-58 leads to the formation of a hybrid FN3_19/22 domain. The top segment of the panel shows the exon structure and corresponding protein domains before exon skipping, while the middle segment displays the structure following exon skipping. The bottom segment highlights the resultant hybrid domain next to the ‘flower’ like protrusion, highlighted in beige as a half circle. **B)** 3D AlphaFold2 models depicting the consequences of exon 54-58 skipping on human (left panel) and zebrafish (right panel) usherin. The visualizations focus on the FN3 domains 17-23, where the segments targeted for skipping are highlighted in yellow. Specifically, these segments include two-thirds of FN3_19, FN3_20 and FN3_21 completely, and one-third of FN3_22. In both human and zebrafish models, the exon skipping results in the creation of a hybrid FN3_19-22 domain. This hybrid domain, annotated in pink, mimics the wild-type protein structure but with a modified configuration. The top half of each panel shows the initial structure before exon skipping, while the bottom half illustrates the resulting hybrid domain structure. **C)** Displays the structural alignment of human Hybrid FN3_19-22 (in red) with human FN3_19 (in gray). The root mean square deviation (RMSD) between these two structures is 0.942 Å. **D)** Displays the structural alignment of human Hybrid FN3_19-22 (in red) with human FN3_22 (in gray). The root mean square deviation (RMSD) between these two structures is 1.67 Å. **E)** Illustrates the alignment of human Hybrid FN3_19-22 (in red) with a consensus structure (in gray). The consensus structure represents an averaged or generalized form derived from multiple alignments of the FN3 domains found in usherin. **F)** Displays the structural alignment of zebrafish Hybrid FN3_19-22 (in red) with FN3_19 (in gray). The root mean square deviation (RMSD) between these two structures is 2.323 Å. **G)** Displays the structural alignment of zebrafish Hybrid FN3_19-22 (in red) with fish FN3_22 (in gray). **H)** Illustrates the alignment of zebrafish Hybrid FN3_19-22 (in red) with a consensus structure (in gray).

Therefore, this *in silico* evidence motivated us to pursue experimental validation in zebrafish to confirm the value of this novel prediction tool based on the formation of hybrid domains to expand the options for therapeutic exon skipping strategies.

### Validating the predictive value of therapeutic exon skipping targets resulting in hybrid FN3 domains using zebrafish as a model

To validate if skipping of *USH2A* exon 54-58 indeed could be considered as a viable future treatment option for *USH2A*-associated retinal disease caused by mutations in exon 54-58, we used zebrafish as a model to investigate the expression and subcellular localization of the usherin^Δexon54-58^ protein. We first employed CRISPR/Cas9 technology to generate a stable zebrafish line from which the genomic region encompassing *ush2a* exons 54-58 was specifically excised. Cas9-sgRNA ribonucleoprotein complexes targeting the genomic region up- and downstream of respectively exon 54 and 58 were injected in single cell-staged fertilized embryos. Anticipated exon excision events were confirmed by genomic PCR and Sanger sequencing (**Figure 4A**). A stable homozygous excision line was bred from a germline-positive founder fish, and was designated *ush2a*^*Δexon54-58*^ or *ush2a*^*rmc41*^. Homozygous *ush2a*^*Δexon54-58*^ fish were viable and did not display any abnormalities in overall body morphology, development, or swimming behavior. To assess the effect of *ush2a* exon 54-58 excision at the transcript level, total RNA was isolated from wild-type and homozygous *ush2a*^*Δexon54-58*^ zebrafish larvae (5 dpf). RT-PCR analysis using forward and reverse primers in exons 50 and 63 of the zebrafish *ush2a* gene, respectively, revealed a shortened amplicon in *ush2a*^*Δexon54-58*^ zebrafish in the absence of any clear alternatively spliced *ush2a* transcripts (**Figure 4B**). Sanger sequencing of the PCR products confirmed the absence of exons 54-58. We next determined whether excision of zebrafish *ush2a* exons 54-58 resulted in the synthesis and correct subcellular localization of usherin^Δexon54-58^ in the retina of homozygous *ush2a*^*Δexon54-58*^ zebrafish larvae. Antibodies directed against the intracellular region of zebrafish usherin were used to label unfixed retinal cryosections of 5 dpf zebrafish larvae (**Figure 4C**). Usherin was previously shown to be present at the photoreceptor periciliary membrane, adjacent to the basal body and connecting cilium marker centrin (13,17). As anticipated, usherin localized to the periciliary region in both wild-type and *ush2a*^*Δexon54-58*^ larvae, and was absent in photoreceptors of *ush2a*^*rmc1*^ knockout larvae. Quantification based on the intensity of the fluorescent signals revealed that usherin^Δexon54-58^ was expressed at at lower levels than wildtype usherin (**Figure 4D**). Adgrv1, a member of the USH2 protein complex, is dependent on the presence of usherin for its localization at the photoreceptor periciliary region, as shown by the significantly reduced Adgrv1 signals in *ush2a*^*rmc1*^ larvae. From another end, the expression levels of Adgrv1 in the *ush2a*^*Δexon54-58*^ larvae is comparable to wildtype (**Figure 5A,B**). This indicates that skipping of exon 54-58 results in the expression of a shortened usherin^Δexon54-58^ protein at a reduced level as compared to wildtype usherin, but that this is sufficient to facilitate the constitution of the USH2 protein complex in zebrafish photoreceptor cells. It was previously shown that the transport of rhodopsin from the photoreceptor inner to outer segment is impaired in zebrafish *ush2a* knock-out models. Despite the slightly reduced levels of usherin expression in the *ush2a*^Δ*exon54-58*^ larvae a normal distribution of rhodopsin is observed, comparable to age- and strainmatched wildtype controls (**Figure 5C,D**), observed in two biological replicates.

**Figure 4.**
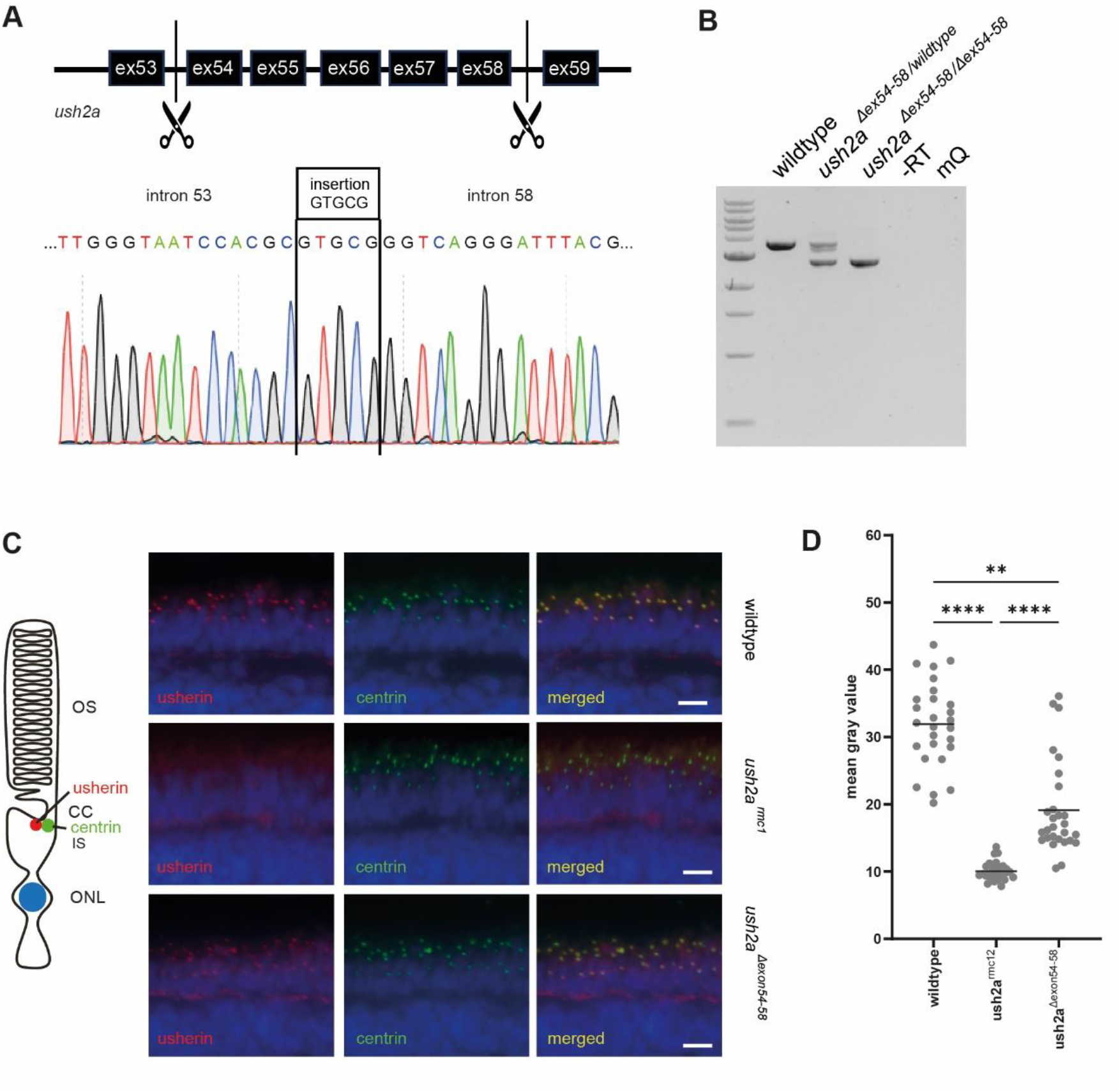
Validation of predicted *ush2a* exon skipping target in zebrafish. **(A)** Using CRISPR/Cas9 genome editing technology, *ush2a* exons 54-58 were successfully excised from the zebrafish genome. Sanger sequencing confirmed excision of the target exons and the simultaneous insertion of 5 nucleotides (GTGCG) during repair. **(B)** Transcript analysis on wildtype, heterozygous and homozygous *ush2a*^*Δexon54-58*^ larvae (5 dpf) revealed the absence of the target exons from the *ush2a* transcript (lower fragment). **(C)** Immunohistochemical analysis showed the expression of usherinΔexon54-58 (red signal) with a subcellular localization in photoreceptor cells, adjacent to basal body and connecting cilium marker centrin (green signal). Nuclei are stained with DAPI (blue signal). Scale bars represent 10 µm. **(D)** Quantification of anti-usherin signal intensity at the periciliary region. Individual data points represent the mean fluorescent intensity of a central single section of a zebrafish eye. Horizontal bar is the mean signal intensity per genotype (n=27-28 eyes). Data were analyzed using one-way ANOVA followed by Kruskal-Wallis multiple comparisons test. ** P < 0.01, **** P < 0.0001.

**Figure 5.**
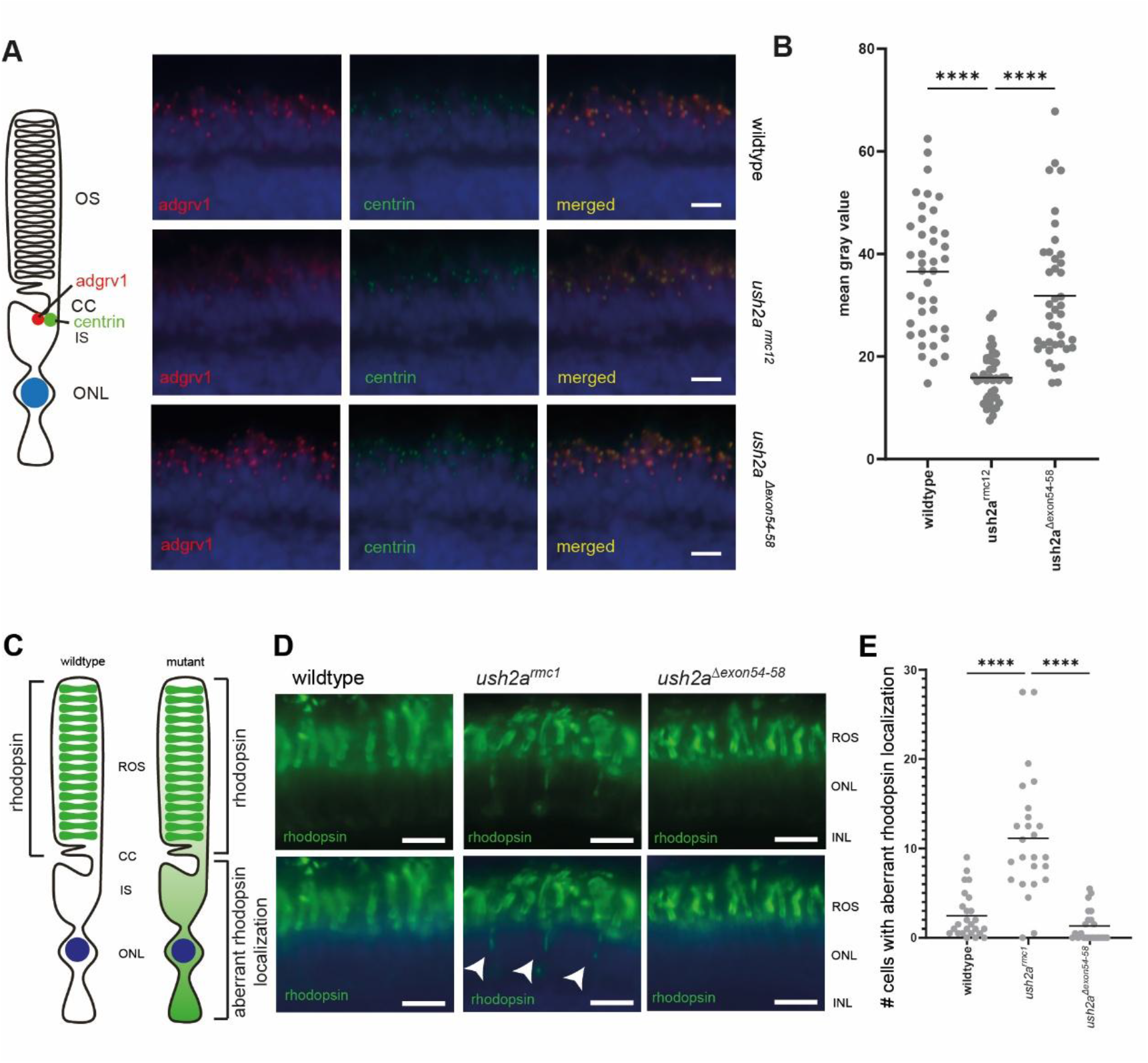
Functional analyses of zebrafish *ush2a*^*Δexon54-58*^ larvae. **(A)** Subcellular localization of Adgrv1 (red) in photoreceptor cells, adjacent to basal body and connecting cilium marker centrin (green). Nuclei are stained with DAPI (blue signal). Scale bars represent 10 µm. **(B)** Quantification of anti-Adgrv1 signal intensity at the periciliary region. Individual data points represent the mean fluorescent intensity of a central single section of a zebrafish eye. Horizontal bar is the mean signal intensity per genotype (n=40-42 eyes). Data were analyzed using one-way ANOVA followed by Kruskal-Wallis multiple comparisons test. **** P <0.0001. **(C)** Schematic representation of a photoreceptor in a wildtype retina, and a photoreceptor with aberrant rhodopsin in a mutant retina. **(D)** Representative images of retina cryosections (6dpf) stained for rhodopsin (green). Nuclei are stained with DAPI (blue). Arrows show aberrant rhodopsin localization. Scale bars represent 10 µm. **(F)** Quantification of number off cells with aberrant rhodopsin localization per genotype (n=22-25 eyes). Data analyzed with one-way ANOVA followed by Kruskal-Wallis multiple comparisons test. **** P <0.0001.

## DISCUSSION

In this study, we explored the added value of integrating next generation *in silico* bioinformatic algorithms to accurately predict protein domain architecture and 3D structural folding into the currently existing preclinical therapeutic exon skipping development pipeline for the future treatment of *USH2A*-associated retinal disease. Our primary focus was on the region encoding the stretch of repetitive FN3 domains within usherin, which harbors a large number of unique (likely) pathogenic variants. By combining existing data on crystallized structures of FN3 domains with deep learning– based protein modeling, we aimed to expand therapeutic opportunities beyond traditional protein domain-based exon skipping approaches by predicting exon skipping events resulting in the formation of correctly folded hybrid FN3 domains, followed by a functional validation using zebrafish models.

Historically, the first clinical applications of exon skipping, such as in Duchenne muscular dystrophy (DMD), relied primarily on sequence-based predictions, without using the benefit of advanced structural modeling (18). At the time, computational tools like AlphaFold2 did not exist, leaving researchers unable to accurately assess the consequences of exon removal on protein structure. As a result, current DMD exon skipping therapies yield shortened proteins that partially rescue the DMD phenotype into a milder “Becker-like” phenotype, rather than restoring full wildtype function (19). The presented hybrid approach differs fundamentally on two levels from the current methods in exon skipping. First, we apply 3D structural modeling and *in silico* analysis before selecting target regions for therapeutic exon skipping. This allows us to get an accurate insight into the 3D architecture of the full length protein and the domain-specific properties. This will increase precision in choosing candidate exons that would yield a functional protein upon skipping. Second, with a detailed 3D overview, one is not restricted to select exons that exactly encode a full domain. Theoretically, almost any exon combination that is in frame and yields a hybrid domain that structurally closely resembles both parental domains, can be considered as a candidate. This hybrid domain approach exemplifies the flexibility in the design of exon skipping therapies, as it allows for the removal of regions containing pathogenic variants while preserving essential protein structure and function. To put it into the perspective, therapeutic skipping of *USH2A* exons 54-58 as presented in this paper, could benefit a significant number of affected individuals, as 188 (likely) pathogenic variants within this target region have been documented based on the LOVD database (www.LOVD.nl/USH2A; visited on February 12, 2026). For instance, the (likely) pathogenic variants p.(Thr3571Met) and p.(Ile3620Thr) in exons 54 and 55 have been reported 13 and 14 times, respectively (20–22). Another example is the p.(Arg3719His) variant in exon 57, which has been reported 33 times as likely pathogenic (23,24). With modern deep learning-based tools such as AlphaFold2, researchers are now also revisiting the DMD protein to identify better exon combinations, a direction that highlights the timeliness and broader relevance of protein modeling in exon skipping research (25).

To evaluate hybrid domains, we used the Root Mean Square Deviation (RMSD) score, which quantifies below 2 Å are generally considered to indicate structures with high similarity which likely have a preserved function. Scores in the range of 2–3 Å suggest moderate similarity, where some deviations are present but may still be tolerated without major loss of function. Values above 3 Å typically reflect poor similarity and carry a greater risk of misfolding and might be indicative for a loss of protein function. However, RMSD values must be interpreted as guidelines rather than absolute thresholds, particularly for domains with flexible regions. For example, in our case study of the zebrafish FN3_19– 22 hybrid, the RMSD exceeded 2 Å due to variations in loop and strand lengths. Upon hybrid formation, some structures tend to fold into longer or shorter strands or loops when compared to the parental domains. To address this, we generated a consensus model based on all FN3 domains, serving as a benchmark to put structural deviations into perspective. While superimposition against the consensus cannot (yet) be quantified numerically, it provides a structural benchmark to decide whether a hybrid matches with the general folding of FN3 domains. This approach balances numerical scoring with structural reasoning, making both RMSD and TM scores useful, but not definitive, tools in hybrid domain functionality prediction.

An important foundation of our work is the observation that domains of the same type generally adopt highly similar folds, even across different proteins. We leverage this fact when predicting the viability of hybrid FN3 domains. Our all-against-all comparison of FN3 domains in usherin shows a shared conserved β-strand architecture, with loops showing variable tolerance to structural deviations. Domains FN3_18 and FN3_19 show strikingly lower TM scores when compared to the other domains. Domain FN3_18 can be identified as special, since this domain seems to act like a scaffold for the ‘flower-like’domain and thereby slightly deviates in its structure. Similarly, domain FN3_19 shows lower TM scores across the comparisons with all other FN3 domains which can be explained by the variation in length and number of its structure elements, even though it consists mostly of a typical FN3 core architecture. Wether the lower expression levels of the FN3_19-22 hybrid protein can be related to structural stability of the new hybrid domain is currently unclear but not unthinkable. The positive results obtained during the validation experiments underline the fact that usherin indeed tolerates variation within this domain and that our approach still leads to a viable exon skip target.

This principle of repeating domain structures is not unique to FN3 domains: proteins such as ADGRV1 contain repetitive calx-β domains that also fold in a conserved structure (26). Recognizing this pattern allows us to extend our predictive framework to other disease-associated proteins with repetitive domain architectures such as cadherins, spectrins, or ADGRV1 (27–29). By understanding which structural elements must be preserved for domain stability and folding, we can systematically design personalized exon skipping strategies based on a patient’s genotype that would result in the formation of correctly folded and therefore functional hybrid domains.

While promising, our pipeline faces important limitations. First, RMSD and TM-align scores are only proxies for structural similarity and cannot alone determine functionality. Experimental validation remains essential. Second, consensus modeling introduces a conceptual scaffold but cannot yet predict to which extend structural deviations, such as unusually long loops or shortened β-strands that have not been seen within the consensus, are functionally tolerated. A central unanswered question is whether only core amino acids within the β-strands are necessary to maintain FN3 domain stability, or if other structural features are also critical. Until these aspects are clarified, the definition of a “functional hybrid domain” remains incomplete. Furthermore, it is important to acknowledge the pitfalls associated with an overreliance on predictive models—including AlphaFold2—without sufficient experimental validation (30). Such reliance can create overconfidence in computational predictions and potentially lead to misinterpretations of protein structure and function. This risk is heightened when analyzing proteins with few structural analogs or those with unique biophysical properties, where model training data are sparse (31). While AlphaFold3 (AF3) introduces important advances, including improved predictions of protein–ligand, protein–nucleic acid, and protein–protein interactions, even state-of-the-art methods can generate plausible yet inaccurate structures in the absence of experimental constraints. To enhance reliability, future pipelines should integrate physics-based approaches such as molecular dynamics (MD) simulations (32), which complement AI-driven predictions and provide additional confidence in structural stability. Looking ahead, advances in AI-based structure prediction (e.g., AlphaFold3 and beyond), combined with MD simulations and wet-lab validation, may allow more reliable *in silico* assessments of hybrid domains. We envision that combining deep learning models with experimental data will lead to a progressively more accurate development pipeline for therapeutic exon skipping.

In conclusion, our work demonstrates that integrating domain knowledge with modern structure prediction tools can overcome some of the inherent limitations of traditional exon skipping approaches. By targeting combinations of exons that create functional hybrid domains, we expand the therapeutic possibilities for *USH2A* and potentially other genes that encode proteins with a repetitive domain architecture. With this work, we aim to broaden the concept of exon skipping, which has traditionally been viewed as a protein-domain oriented approach combined with the distribution of (recurrent) pathogenic variants, based on 2D structural predictions (SMART database). Our hybrid approach based on accurate 3D structural modelling shifts this paradigm by enabling the removal of mutational hot spots on a larger scale than was previously possible. While challenges remain in defining and validating functional hybrids, this research lays the foundation for a new era in exon skipping, one that combines computational precision with translational potential to design more effective therapies.

## MATERIALS AND METHODS

### usherin SMART and UniProt models

The two-dimensional domain structure of the usherin protein product was detailed using the SMART (16) and UniProt (33) databases. The Swissprot sequence O75445 was used as an input to search the SMART and UniProt databases to obtain a comprehensive two-dimensional domain overview. This process facilitated the detailed visualization of domain architectures and identification of potential functional sites within the protein.

### 3D structural modeling of usherin using AlphaFold2

The sequence was divided into segments of 300 amino acids, with each batch overlapping the previous one by 7 amino acids. Each segment of the complete human usherin protein was modeled using the AlphaFold2 Google Colab notebook (34). The 3D overview of the full protein was constructed by visualizing each Alphafold2 model with the PyMol 3.1 visualization program (35).

### Exon-to-domain boundaries identification

The Human Genome Browser was used to compile a list of all usherin encoding-exons (36). These exons were then mapped directly onto the identified protein domains using PyMol to establish an exon-to-domain boundaries overview. This network facilitated the improved selection of exon candidates.

### Selection of exon skipping targets and modeling of hybrid domains

Potential exons that encode the stretch of fibronectin type 3 (FN3) domains that contain the highest frequency of pathogenic variants were identified using the LOVD database (4). In frame exon combinations resulting in the formation of a hybrid FN3 domain were determined. The sequences of selected exons were removed from the coding sequence and the remaining transcript was subsequently translated into protein, prior to modeling. Hybrid AlphaFold2 modeling was conducted in segments of three predicted domains, with a hybrid domain positioned in the middle of the chain, to verify the feasibility of chain formation.

### Zebrafish husbandry and maintenance

All animal experiments were conducted in accordance with the Dutch guidelines for the care and use of laboratory animals (Wet op de Dierproeven 1996) and European regulations (Directive 86/609/EEC), as approved by the Dutch Ethics committee of the Central Committee Animal Experimentation (Centrale Commissie Dierproeven [CCD]; application number AVD10300 2022 15892). Wild-type Tupfel Longfin (TL) zebrafish were used. Zebrafish were maintained and raised according to standard methods (37). Both zebrafish larvae and adults were kept at a 14 hours of light and 10 hours of darkness regime. Adult zebrafish were fed once a day with Gemma Micro 300 dry pellets (#13177, Zebcare, Nederweert, The Netherlands) at an amount of 5% of their average body weight and once a day with Artemia sp. nauplii. Zebrafish embryos were obtained from natural spawning.

### CRISPR/Cas9 genome-editing design

For the generation of the *ush2a*^*Δexon54-58*^ zebrafish, single guide RNAs (sgRNAs) targeting zebrafish *ush2a* introns 53 and 58 (NCBI accession XM_009293147.3)(**table S1**) (38) were identified with the online web tool CHOPCHOP (https://chopchop.cbu.uib.no/). sgRNAs for which no off-target sites were predicted and which had the highest predicted efficiency score were selected and ordered (Integrated DNA Technologies, Coralville, USA) (39). 5’ sgRNA (50ng/µl), 3’sgRNA (50ng/µl) and commercial Alt-R S.p. Cas9 Nuclease V3 (#1081059, IDT, Newark,NJ, USA)(800ng/µl) were injected as discribed previously (15). The *ush2a*^*rmc1*^ mutant was described previously (17).

### Genotyping

Genomic DNA was extracted from whole larvae (1 day post fertilization (dpf)) or caudal fin tissue from adult zebrafish. Tissue was lysed in 25 μl (larvae) or 75 μl (fin tissue) lysis buffer (40 mM NaOH, 0.2 mM EDTA) at 95°C for 20 minutes. The lysed samples were neutralized with 10% (v/v) 1M TRIS-HCl (pH 7.5) and diluted 10 times with Milli-Q water. 1 μl of diluted sample was used as a template in PCR to amplify the zebrafish *ush2a*^*Δexon54-58*^ allele or the *ush2a*^*rmc1*^ allele using standard cycling conditions. All primer sequences are listed in **Table S2**. The presence of the introduced lesions and the subsequent presence or absence of the zebrafish *ush2a*^*Δexon54-58*^ or the *ush2a*^*rmc1*^ allele in the following generations was confirmed by Sanger sequencing.

### RNA isolation, cDNA synthesis and transcript analysis

Total RNA from zebrafish larvae (5 dpf) was extracted using the RNeasy® Micro kit (#74004, Qiagen, Hilden, Germany), according to manufacturer’s instructions. cDNA was synthesized using SuperScript™ IV Reverse Transcriptase (#18090010, Thermo Fisher Scientific,Waltham, MA, USA), combined with an oligo(dT) primer (#18418012, Thermo Fisher Scientific, Waltham, MA, USA), according to manufacturer’s protocol.The target region of *ush2a* exon 50 till 63 was amplified from the synthesized zebrafish cDNA using Q5® High-Fidelity DNA Polymerase (#M0491L, New England Biolabs, Ipswich, MA, USA). Primer sequences are listed in **Table S2**.

### Immunohistochemistry on zebrafish cryosections

Immunohistochemistry on 5 dpf or 6dpf zebrafish larvae and subsequent quantification of the fluorescent signals has been performed as previously described (13). The following primary antibody and dilution was used to assess usherin or Adgrv1 expression and rhodopsin localization: rabbit anti-usherin (1:500; #DZ01481, Boster Bio, Pleasanton, CA, USA), rabbit anti-Adgrv1 (1:200; #DZ41032; Boster Bio), mouse anti-centrin (1:500; #04-1624, Millipore, Burlington, MA, USA), mouse anti-rhodopsin (1:2000, Clone 4D2; NBP2-59690, Novus Biological, Centennial (CO), USA). Secondary antibodies Alexa Fluor 568 goat anti-rabbit (#A11011, Thermo Fisher Scientific, Waltham, MA, USA) and Alexa Fluor 488 goat anti-mouse (#A11209, Thermo Fisher Scientific, Waltham, MA, USA) were used in a 1:800 dilution. Nuclei were counterstained with DAPI (1:8000; #D1306; Thermo Fisher, Waltham, MA, USA). Images were taken using a Zeiss Axio Imager fluorescence microscope equipped with an AxioCam MRC5 camera (Zeiss, Jena, Germany).

### Statistical analysis

Graphpad Prism 10 software (http://www.graphpad.com/scientific-software/prism/) was employed to generate scatter plots, to calculate mean values, and to perform statistical tests. Differences among groups were assessed using a Kruskal-Wallis test p ≤ 0.05 was considered statistically significant (* p ≤ 0.05, ** p ≤ 0.01, *** p ≤ 0.001, and **** p ≤ 0.0001). n indicates the number of biological replicates, all bars within the graphs represent mean values.

## DATA AVAILABILITY STATEMENT

Data are available from authors upon request

## ACKNOWLEDGEMENTS

This study was financially supported by the Dutch Usher syndrome Foundation (to H.V. and E.v.W.); the Reggeborgh Foundation (R0006760 to E.v.W. and E.d.V.); the Gelderse Blindenstichting (E.v.W.); the Ministry of Education, Culture and Science of the Netherlands (grant: Gravitation 024.006.034 to E.v.W.); Zeldzame Ziekten Fonds (A25-1500; to H.V. and E.v.W.); LSBS, ANVVB, Oogfonds (UZ 2025-8; to H.V. and E.v.W.); Stichting Blindenhulp (2024-59-4521); Rotterdamse Stichting Blindenbelangen (B20240035; to H.V. and E.v.W.); and Dowilvo (2024-192; to H.V. and E.v.W.). D.R. acknowledges the financial support received from the Hypatia Fellowship at Radboudumc (Rv819.52706). We are also grateful to Prof. Dr. Martijn Huijnen and Dr. Dario Marzella for their engaging discussions and valuable advice and acknowledge Antoon van der Horst, Jeroen Boerrigters and Kimberly Janssen from the Radboud University Zebrafish Facility (www.ru.nl/zebrafish) for their excellent fish husbandry.

## AUTHOR CONTRIBUTIONS

L.M. developed the hybrid method as the core of the study’s conception and design, conducted/supervised the *in silico* research, prepared the initial draft of the manuscript, and created all figures included in the article, except for Figure 4 and 5. D.R. provided daily supervision to L.M. throughout the research process and participated significantly in writing and editing the manuscript. A.L. contributed to the generation of AlphaFold2 3D models and the annotation of exon to domain boundaries. S.B., E.d.V. and T.P. performed the wetlab experiments in zebrafish, while E.v.W. supervised these activities. H.V., as principal investigator, was responsible for the final review of the paper and general supervision of L.M. PACtH critically reviewed the results, their interpretation, and the manuscript. All authors critically reviewed the article.

## SUPPLEMENTARY DATA

**Table S1.**
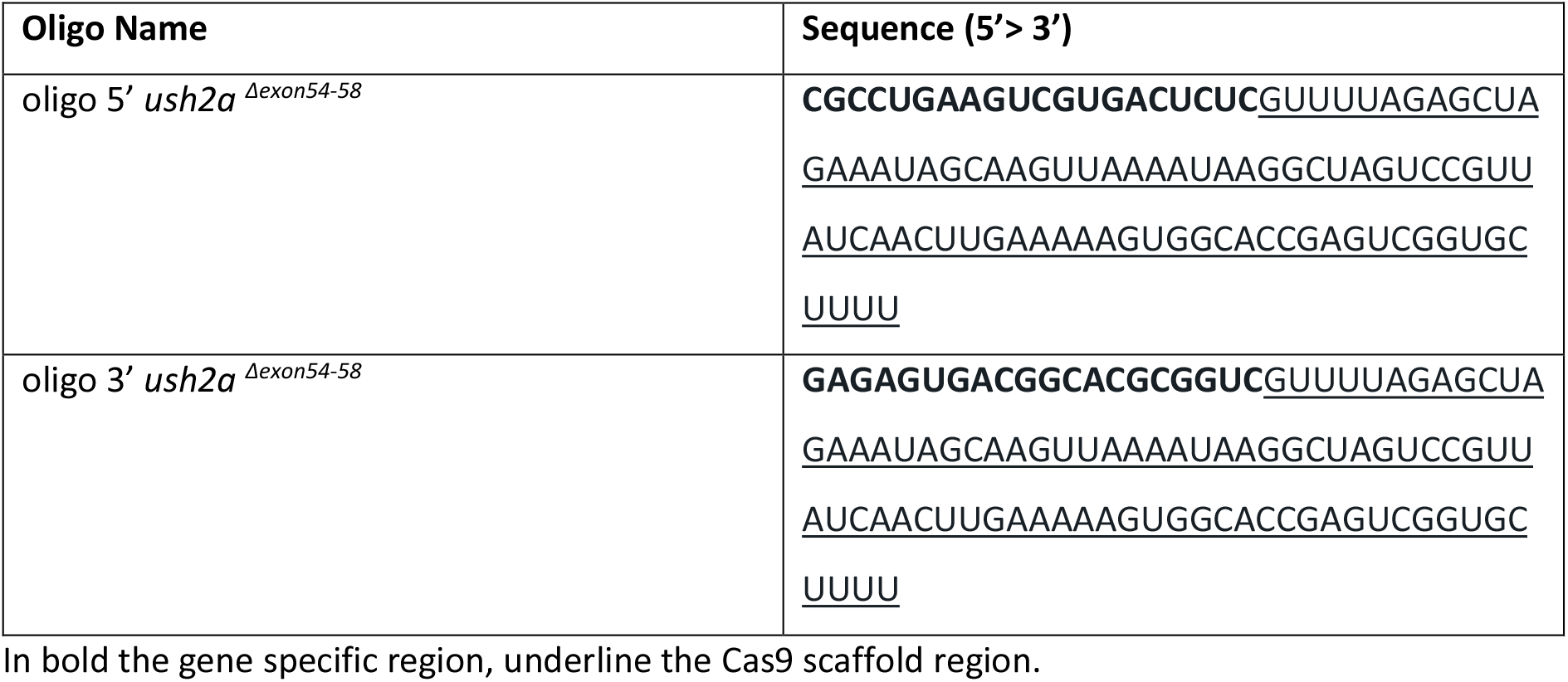
sgRNA used to genetrate *ush2a*^*Δexon54-58*^ zebrafish.

**Table S2.**
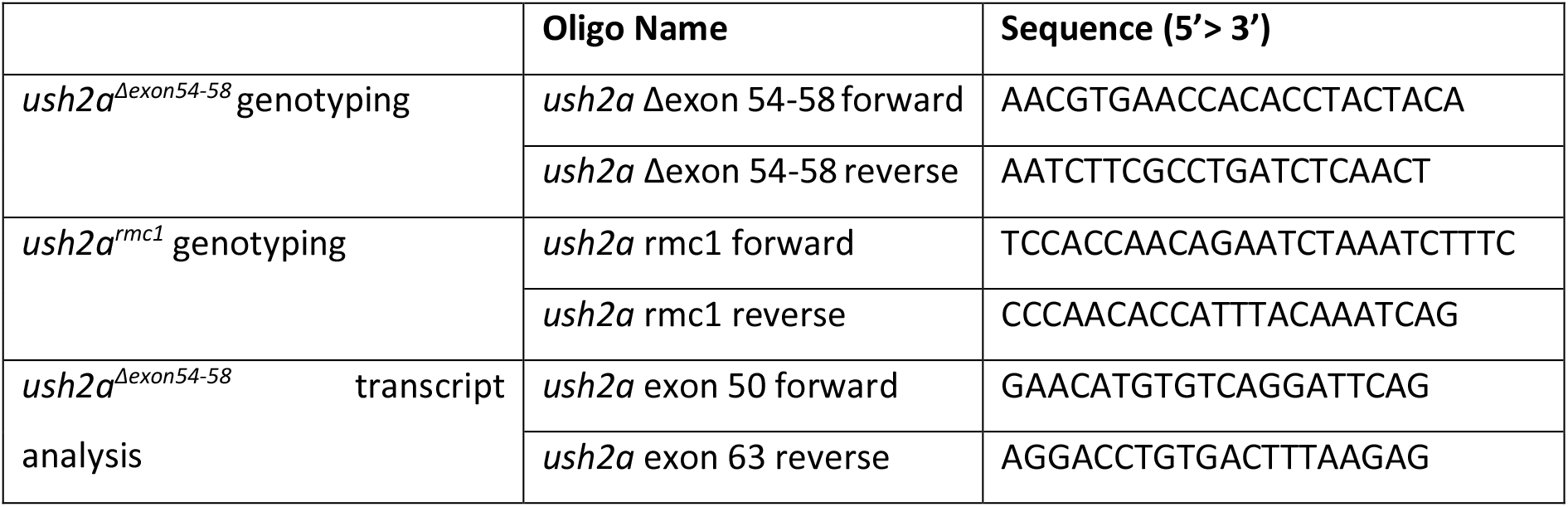
Primers used for genomic and RT-PCR.

**Figure S1.**
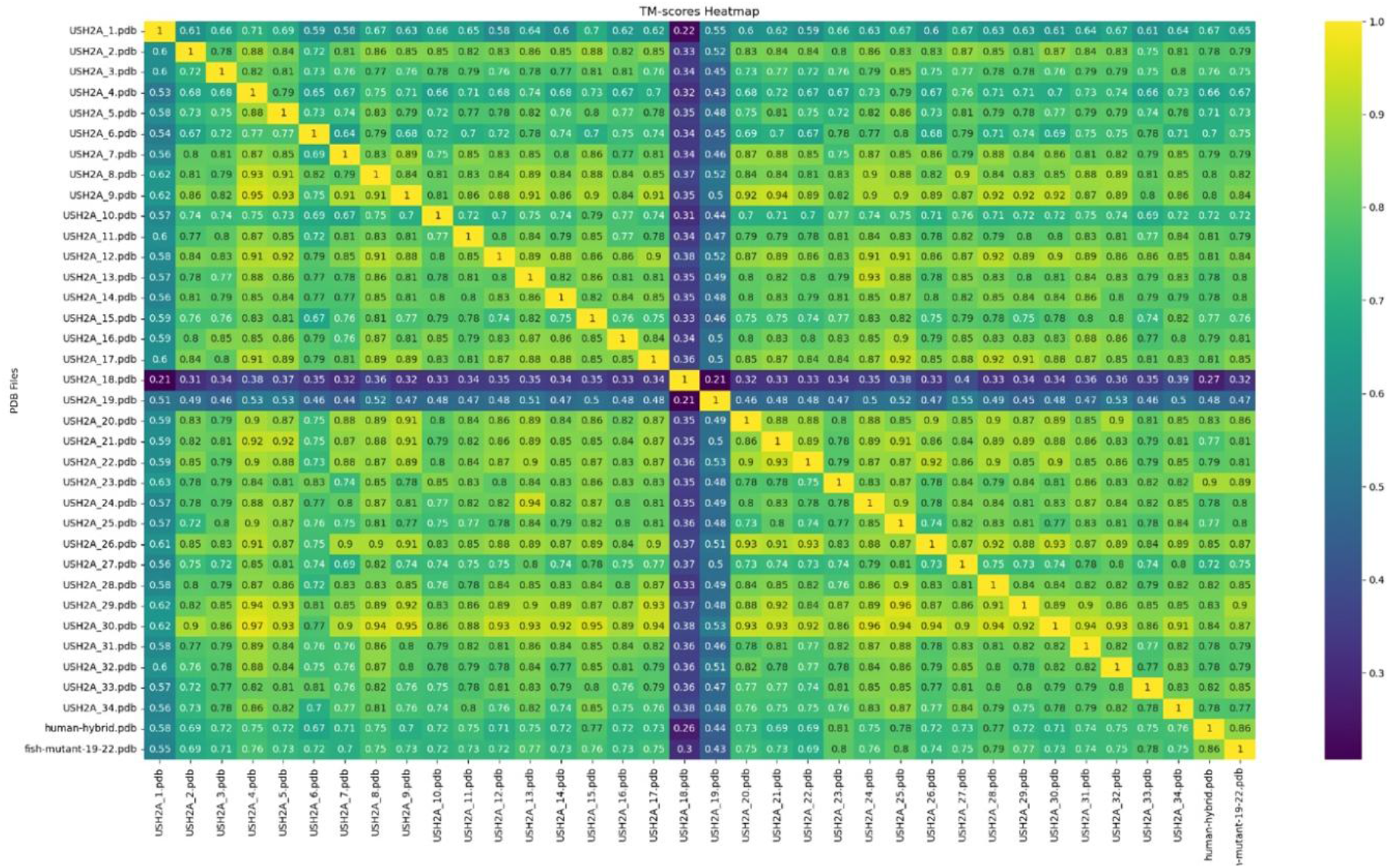
Heatmap of TM-scores for structural similarity among fibronectin type III domains in usherin. A heatmap illustrating the structural similarities between various fibronectin domains within the usherin protein, as measured by TM-scores. Each row and column corresponds to a specific fibronectin domain labeled accordingly (e.g., “USH2A_1_pdb”). The diagonal of the matrix, which is uniformly yellow, represents a perfect TM-score of 1, indicating identical structures when a domain is compared to itself. Off-diagonal values show the TM-scores between different domains, with colors ranging from yellow (high similarity) to blue/purple (low similarity). The heatmap also includes comparisons to hybrid and mutant forms, such as “human-hybrid.pdb” and “fish-mutant-19-22.pdb”, allowing for a comparative analysis of structural variations within these altered structures.

